# Multi-Wavelength Analytical Ultracentrifugation of Biopolymer Mixtures and Interactions

**DOI:** 10.1101/2021.12.29.474408

**Authors:** Amy Henrickson, Gary E. Gorbet, Alexey Savelyev, Minji Kim, Sarah K. Schultz, Xiaozhe Ding, Jason Hargreaves, Viviana Gradinaru, Ute Kothe, Borries Demeler

## Abstract

Multi-wavelength analytical ultracentrifugation (MW-AUC) is a recent development made possible by new analytical ultracentrifuge optical systems. MW-AUC is suitable for a wide range of applications and biopolymer systems and is poised to become an essential tool to characterize macromolecular interactions. It adds an orthogonal spectral dimension to the traditional hydrodynamic characterization by exploiting unique chromophores in analyte mixtures that may or may not interact. Here we illustrate the utility of MW-AUC for representative classes of challenging biopolymer systems, including interactions between mixtures of different sized proteins with small molecules, mixtures of loaded and empty viral AAV capsids contaminated with free DNA, and mixtures of different proteins, where some have identical hydrodynamic properties, all of which are difficult to resolve with traditional AUC methods. We explain the improvement in resolution and information content obtained by this technique compared to traditional single- or dual-wavelength approaches. We discuss experimental design considerations and limitations of the method, and address the advantages and disadvantages of the two MW optical systems available today, and the differences in data analysis strategies between the two systems.

## Introduction

In 2008 the Cölfen lab introduced the first fiber-based UV-visible multi-wavelength detector for the analytical ultracentrifuge [1], adding an optical characterization dimension to the traditional hydrodynamic separation. This accomplishment added an important method to the toolkit of analytical ultracentrifugation (AUC), further enhancing the potential for discovery through the already capable and time-honored method. This optical system was further improved in 2015 [2], and our laboratory contributed the data analysis framework implemented in UltraScan [3] for data generated by this detector [4]. In 2018 this method was further enhanced by the addition of mirror optics [5] (referred to here as “*Cölfen optics*”). This design has been successfully employed in multi-wavelength experiments of biopolymers with chromophores in the visible range [6], protein-DNA mixtures [4], and protein-RNA interactions [7]. The Cölfen optics design has been made available under an open source license [8]; it is intended to be retrofit into a preparative ultracentrifuge sold by Beckman-Coulter. In 2016, Beckman-Coulter released a new generation of analytical ultracentrifuges, the Optima AUC^™^ series. It was equipped with Rayleigh interference optics and multi-wavelength capable UV/visible absorption optics (referred to here as “*Beckman optics*”), and is currently the only commercially available analytical ultracentrifuge. Multi-wavelength experiments with biopolymers performed with the Beckman optics are starting to emerge and include studies on heme proteins [9, 10, 11], triphenylmethane dyes binding to peptide trimers derived from amyloid-β peptides [12], and protein-DNA interactions [13].

### Principles of MW-AUC

Analytical ultracentrifugation is a technique used to measure the partial concentrations, sedimentation coefficients, and the diffusion coefficients of analytes present in colloidal molecular mixtures. From this information, details about the analyte’s size and anisotropy can be obtained [14]. Detection of the molecules is traditionally performed by scanning the sedimenting sample using single-wavelength absorbance spectroscopy as a function of radius and time. In a MW-AUC experiment, the sedimenting sample is scanned at multiple wavelengths. If the solution contains different analytes, each characterized by a different absorbance spectrum, MW-AUC detection provides a second, orthogonal characterization method by resolving analytes not just by differences in their hydrodynamic properties, but also by their absorbance spectra. If the intrinsic molar extinction profiles for each pure analyte are known, and they are sufficiently dissimilar, the spectrum of the mixture can be decomposed into the partial absorbance contributions from each analyte, and the molar quantity of each constituent can be determined [7]. Molecules that form complexes will sediment faster than their unbound forms due to the increase in mass of the complex. The stoichiometry and molar ratio of each analyte in the complex can be deduced by integrating the decomposed spectra. This second dimension adds important information to the hydrodynamic properties, extending the value and impact of traditional AUC.

### Differences between the two MW-AUC optical systems

While both UV/visible optical systems mentioned above share mirror-based optics and support the acquisition of experimental data at multiple wavelengths, important differences in the two systems affect how data are collected, stored, and analyzed. These differences also determine the types of experiments that can be performed with the instruments. Both optics use a stepping motor to scan the radial domain rapidly. The Cölfen optics employ a data collection system where white light passes through the sample, and then through an optical fiber to a diffraction grating. The diffracted light is then imaged on a linear CCD spectrophotometer, producing a wavelength intensity scan with approximately 0.5 nm resolution for each radial position imaged with the device. In the Beckman optics, white light passes over a diffraction grating before passing monochromatic light with 1 nm resolution through the sample. The resulting monochromatic intensity is imaged for each wavelength sequentially on a photomultiplier tube at each radial position in the AUC cell, producing multiple single-wavelength velocity experimental data sets.

### Advantages and limitations for each MW-AUC optical system

These fundamentally different optical systems have pros and cons to be considered in the design and analysis of experiments. The most significant difference between the different optical designs is the order in which data are collected. With the Cölfen optics, experimental data from different wavelengths are collected in parallel, which offers a distinct scanning speed advantage. The Beckman optics employ a photomultiplier tube which scans monochromatic light at a single wavelength, requiring each wavelength to be acquired sequentially. The use of a photomultiplier tube offers distinct dynamic range advantages, especially in the lower UV range, where fiber-based CCD systems suffer from reduced light intensity and therefore lack sufficient sensitivity. This presents a problem for the case of biopolymers (nucleic acids, proteins, lipids, carbohydrates), where detection often relies on the measurement of chromophores that absorb between 210 nm – 240 nm (see Figure SI 1). This lack of sensitivity is further amplified when buffer components that absorb below 260 nm are used, because it decreases the dynamic range available for the detection of the intended analytes. Higher sensitivity can be achieved with the photomultiplier design by scaling the photomultiplier voltage, and therefore, for measurements below 240 nm the Beckman optics are preferred. On the other hand, serial wavelength detection imposes significant throughput limitations, especially when more than 20 wavelengths, or more than two samples need to be measured in a single run. Since the Cölfen optics permit the simultaneous acquisition of a broad range of closely spaced wavelengths for multiple cells, these optics are eminently well suited for measuring systems where chromophores need to be examined over a large wavelength range, especially in the visible range where the Cölfen optics have sufficient dynamic range. When using UltraScan to acquire multi-wavelength data from the Beckman optics [15], data acquisition is restricted to a maximum of 100 wavelengths per cell, but they do not have to be spaced in regular intervals. However, 100 wavelengths are often too many, especially for rapidly sedimenting analytes, since significant delays are encountered during the initial calibration of the photomultiplier gain setting, which needs to be performed for each wavelength and channel. This delay prevents data collection at the beginning of the experiment, causing potential loss of detection for large molecules and aggregates. Consequently, the scan frequency for each wavelength is significantly decreased, despite rapid radial scanning. For experiments with more than 15-20 wavelengths, it is often not advisable to scan more than a single cell, while in the Cölfen optics, all rotor positions can be filled, and scans early in the experiment are not missed. With the Beckman optics, it may be necessary to sacrifice sedimentation resolution by scanning rapidly sedimenting analytes at slower than optimal rotor speed to gain more time for scanning. Nevertheless, the signal-to-noise level in the Beckman optics is exceptional, typically resulting in residual mean square deviations of less than 2.0 x 10^-3^ absorbance units, with a radial resolution of 0.001 cm. Hence, comparable statistics can be achieved with the Optima AUC even with fewer scans in a sedimentation velocity experiment.

## Results

The hydrodynamic separation of free and associated analytes alone often does not provide sufficient resolution to permit a clear and unambiguous interpretation of AUC experiments for two important reasons: First, different analytes may have similar hydrodynamic properties, such as size, anisotropy, and density, and therefore would not be distinguishable by hydrodynamic separation. Secondly, the ability to uniquely identify each analyte decreases with an increasing number of analytes because the observed signal is proportional to the relative amount of each analyte. If too many analytes are present, it is impossible to distinguish them based on hydrodynamic information alone. In MW-AUC experiments, the additional spectral information provides a second dimension to identify analytes by their unique chromophores. We distinguish two basic experiments: a) cases where spectral properties are not available in pure form for each unique chemical species present in a mixture, and b) cases where the pure spectra for each unique chemical species are known, and molar extinction coefficients are available for each measured wavelength. In the case of (a), it is still possible to extract and review the spectral properties after hydrodynamically separating all species. Even though molar extinction coefficients may not be available, the spectral pattern may still provide useful insights. For cases described by (b), a mathematical deconvolution of spectral contributors will then identify the chemical nature of each hydrodynamic species, and for complexes, the stoichiometry of assembly. Examples for both cases are discussed below. In a MW-AUC experiment, multiple datasets from traditional single-wavelength experiments are collected at multiple wavelengths and combined for a global analysis, which can extract a second approach to characterize the identity of the analytes, based on their unique spectral contributions to the overall signal. Since different types of biopolymers have unique spectral properties, it is therefore possible to resolve them not only based on their hydrodynamic properties, but also based on their unique spectral properties.

### a) Hydrodynamic separation of spectral components

In cases where absorbance spectra from individual analytes with unique spectral characteristics cannot be obtained in pure form for all components in a mixture, an optical deconvolution of individual analytes will not be possible. Instead, a different strategy can prove valuable. It displays the spectral profiles of the hydrodynamically separated species. This approach can be very effective and useful, provided multiple components in the mixture can be hydrodynamically separated. A representative example of this approach was demonstrated for mixtures of CdTe quantum dots by Karabudak et al. [16], where 24 unique hydrodynamic species were identified, and unique spectral properties of at least seven components could be derived over the examined wavelength range. In this method, s-values with non-zero amplitude obtained at different wavelengths are integrated at each wavelength to generate a spectral absorbance pattern for each unique hydrodynamic species.

The hydrodynamic separation of biopolymers typically has a lower resolution than the highly dense metal quantum dots. However, if hydrodynamic separation is achieved, this method is still effective for classifying individual components. UltraScan offers a three-dimensional (3D) viewer, which projects the integrated sedimentation profile as a function of wavelength.

#### Example 1 Identification of components in an oil seed protein extract

Figure 1 shows the MW-AUC results for a heterogeneous oil seed plant protein extract after removing the lipid phase. In this example, the plant extract contained polyphenols, small molecules with a 315 nm absorbance peak, and proteins. A MW-AUC experiment was successfully able to answer the following questions: 1. Are the polyphenols free in solution or bound to the protein? 2. Are the proteins degraded or intact? First, the polyphenols, identified by their 315 nm absorbance peak, sedimented as expected with a very low sedimentation coefficient (~1S) and did not appear to be bound to any larger molecules. Furthermore, a peak around 12S displayed a spectral signature of a typical protein with an absorbance maximum around 280 nm. The protein sedimenting at 12S has a narrow distribution suggesting that this protein is intact, and the absence of 315 nm absorbance indicates that no polyphenols are bound. A smaller amount of protein signal was found at < 2S, suggesting the presence of a second protein or a small fraction of a potentially degraded protein. Its absorbance spectrum also suggests the absence of any polyphenols binding to it.

**Figure 1:**
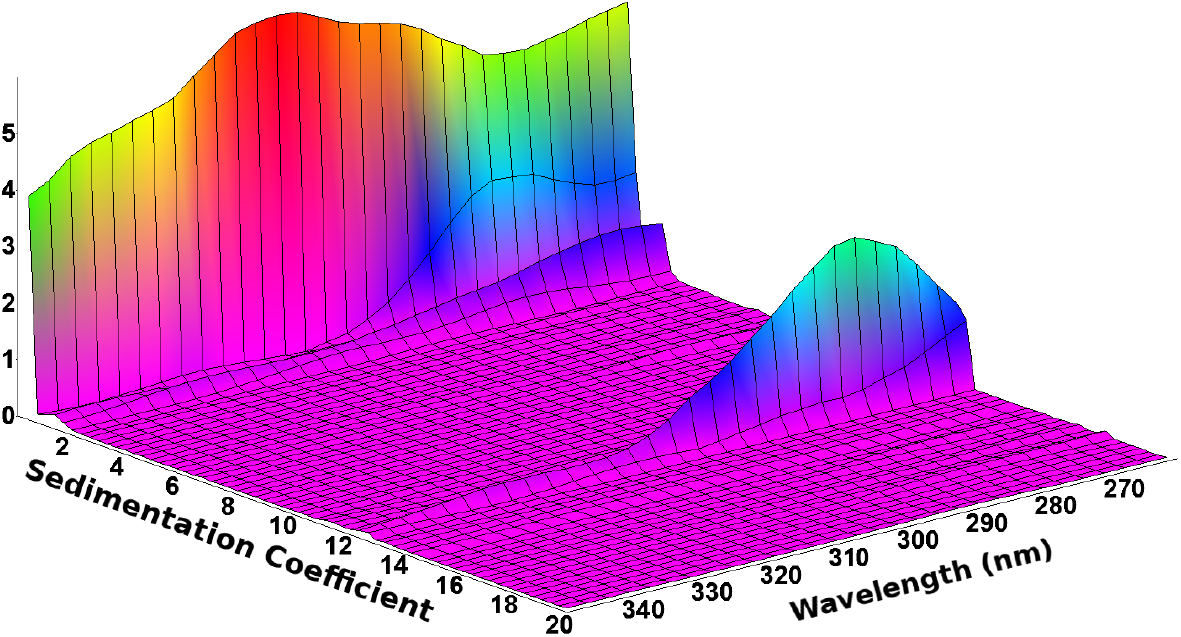
MW-AUC sedimentation velocity experiment of oil seed protein extracts. Polyphenols are small molecules not associated with larger proteins and absorb with a maximum at 315 nm (~1S), while protein peaks absorb with a maximum at 280 nm (~12S). Data collected with the Beckman Optics.

### b) Spectral separation of hydrodynamic components

If pure spectra are available for individual species in a mixture, along with their molar extinction coefficients, spectral decomposition can be applied to determine absolute molar amounts of each species, whether free in solution or interacting with another molecule. In this case, also the stoichiometry of interaction is available. A large class of experimental applications lend themselves to this approach.

#### Example 2 use of fluorescent tags

To study biopolymers without distinct chromophores (lipids, carbohydrates) or protein-protein interactions among proteins with very similar absorbance profiles in the ultraviolet, fluorescent tags or fluorescent protein fusions can be used to impart a unique chromophore to a molecule. Excitation spectra from commercially available fluorescent dyes for tagging biopolymers and fluorescent proteins span a wide range of the visible spectrum and can be used to add a unique chromophore to a molecule of interest. To validate the method, we mixed ultramarine [17], mTeal [18], and mPapaya [19] fluorescent proteins at different ratios and measured their sedimentation between 400-600 nm, the region containing the most significant difference in their absorbance spectra (see SI 2). After spectrally deconvoluting the MW-AUC experimental data, all three species can be baseline-resolved and accurately quantified (see Figure 2). The varying ratios of concentration recovered from the peak integrations shown in Figure 2 accurately reflect the pipetted ratios. The result is even more remarkable, considering that mTeal and mPapaya have identical hydrodynamic properties and would not be distinguishable if measured using traditional single-wavelength AUC. Unlike the monomeric mTeal and mPapaya, Ultramarine sediments as a constitutive dimer with a higher sedimentation coefficient, allowing it to be hydrodynamically separated.

**Figure 2:**
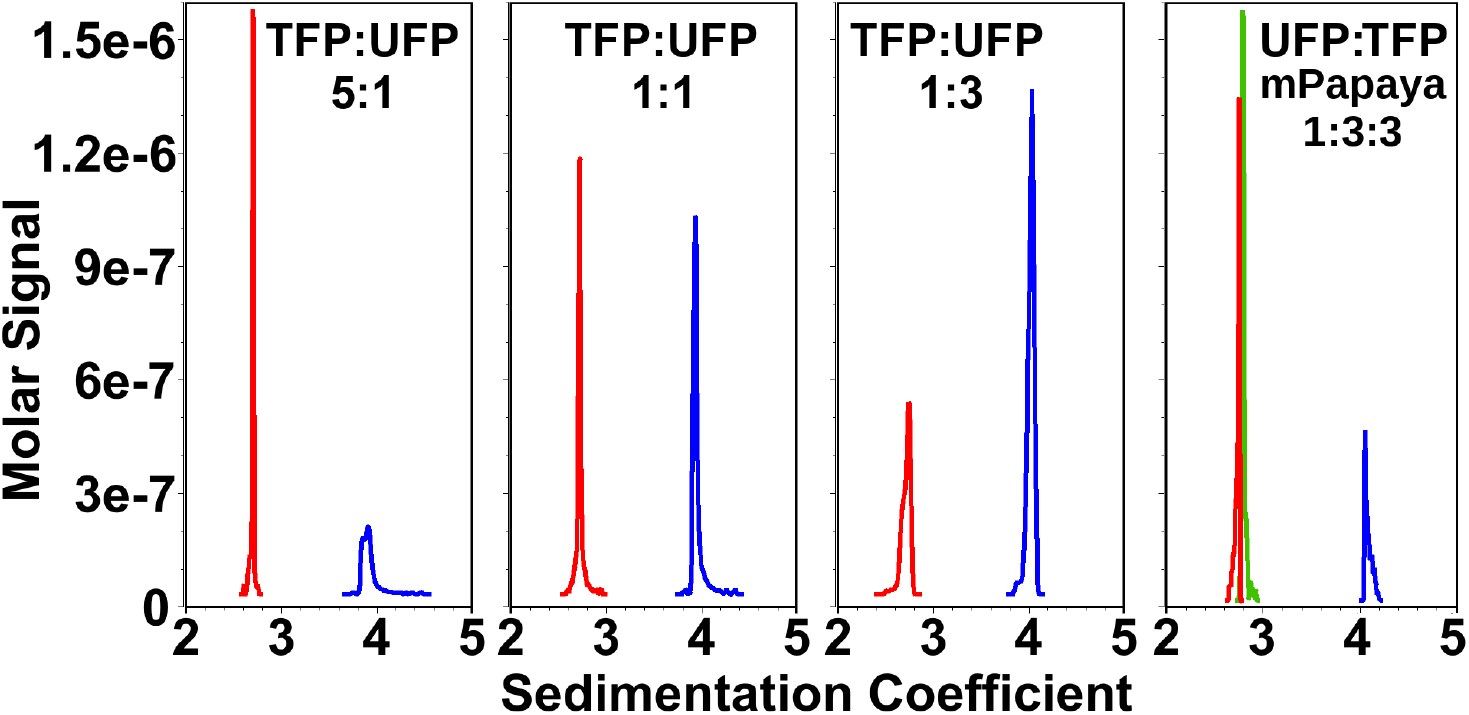
Baseline-resolved mixtures of fluorescent proteins. MW-AUC analysis of mixtures of two or three fluorescent proteins mixed at different ratios: Ultramarine fluorescent protein (UFP, blue), mTeal fluorescent protein (TFP, red), and mPapaya (green). Relative ratios of mixed proteins can be resolved within pipetting error. **Left:** TFP:UFP 5:1, **Center Left:** TFP:UFP 1:1, **Center Right:** TFP:UFP 1:3, **Right:** UFP:TFP:mPapaya 1:3:3. All proteins have identical monomeric mass, but ultramarine fluorescent protein exists as a constitutive dimer which results in a higher sedimentation coefficient. Note that mPapaya and TFP are baseline resolved despite having identical sedimentation coefficients.

#### Example 3 Accurate characterization of viral vector cargo loading

Adeno-associated virus (AAV) formulations deliver genetically encoded tools to cells for gene therapy applications. For these formulations, it is imperative to quantify the nucleic acid cargo loaded into the AAV capsid and to correctly quantify the amounts of empty, partially filled, and full capsids, as well as any contaminants, such as free DNA and protein aggregates [20, 21]. MW-AUC analysis is ideally suited to provide a significantly more realistic insight into viral particle loading than traditional AUC methods which only measure 260/280 nm absorbance [22], or single-wavelength and interference detection [23]. As shown previously, MW-AUC achieves reliable and quantitative separation between protein and DNA signals [4, 24, 25], and is ideally suited to accurately quantify the DNA loading state of AAV capsids. After the spectral deconvolution into the pure DNA and the capsid protein absorption profiles, precise molar ratios of protein:DNA can be assigned to each hydrodynamic species detected in a mixture. By tracking the hydrodynamic signals from protein and DNA separately, the relative amount of capsid protein and DNA in each hydrodynamic species can be readily obtained, and the true ratio of empty, partially loaded, and full capsids can be unambiguously determined. In addition, an assignment of the chemical identity of other peaks not readily assigned to empty, partial, or full capsids is also possible. Figure 3 shows a purified recombinant AAV9 sample analyzed by MW-AUC sedimentation velocity experiments, measuring 230-300 nm, and by single wavelength AUC at 280 and 260 nm for comparison with traditional measurement approaches [24]. The resulting MW-AUC data were deconvoluted into protein (green) and DNA (red) absorbance spectra. These data illustrate several key advantages of the MW-AUC approach. First, the presumed empty capsid species sedimenting at ~65S co-sediments with ~15% of the total DNA signal (red trace), identifying it as either a partially loaded capsid or a capsid with nucleic acid attached to the outside. Second, the ratio of partially filled and completely filled capsid is close to 9:1 (green trace), not 6.5:3 as suggested by the 280 nm single wavelength experiment (cyan trace), or 1:1 as suggested by the 260 nm experiment (blue trace), highlighting the improve resolution. Both 260 nm and 280 nm single wavelength analyses overestimate the filled capsid proportion because of the significant absorbance of DNA at 280 nm, which results in improper interpretations of AAV loading efficiencies. Third, the contaminant sedimenting between 5S-20S is solely composed of nucleic acid and does not contain any protein component, which can not be determined from 260 nm and 280 nm analysis alone. A negative stain transmission electron micrograph (TEM) of the same sample is shown in Figure 4, and illustrates the limitations in resolution when TEM is used for characterization. Not only are contaminating DNA molecules not readily apparent in the TEM, but also empty and partially filled capsids cannot be distinguished. Furthermore, bulk observation in AUC provides improved statistics over single-particle counting methods.

**Figure 3:**
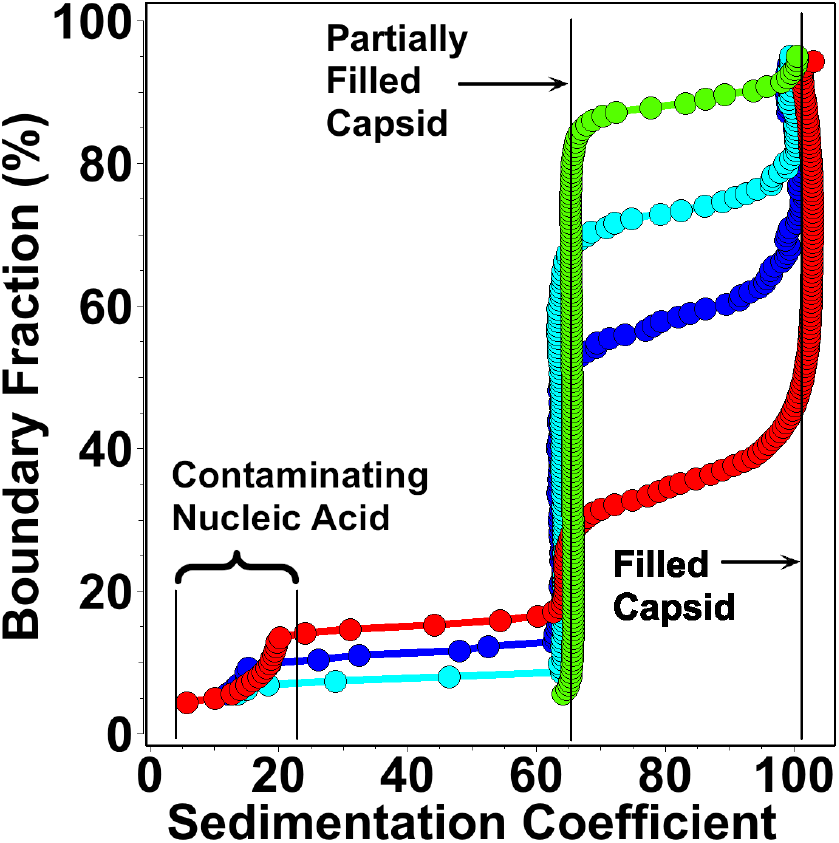
G(s) distributions showing loaded (~100S) and partially filled (~65S) AAV particles, and nucleic acid contaminants. Experiments were performed at 260 nm (blue), 280 nm (cyan), and with MW-AUC deconvoluting protein signal (green) from DNA signal (red).

**Figure 4:**
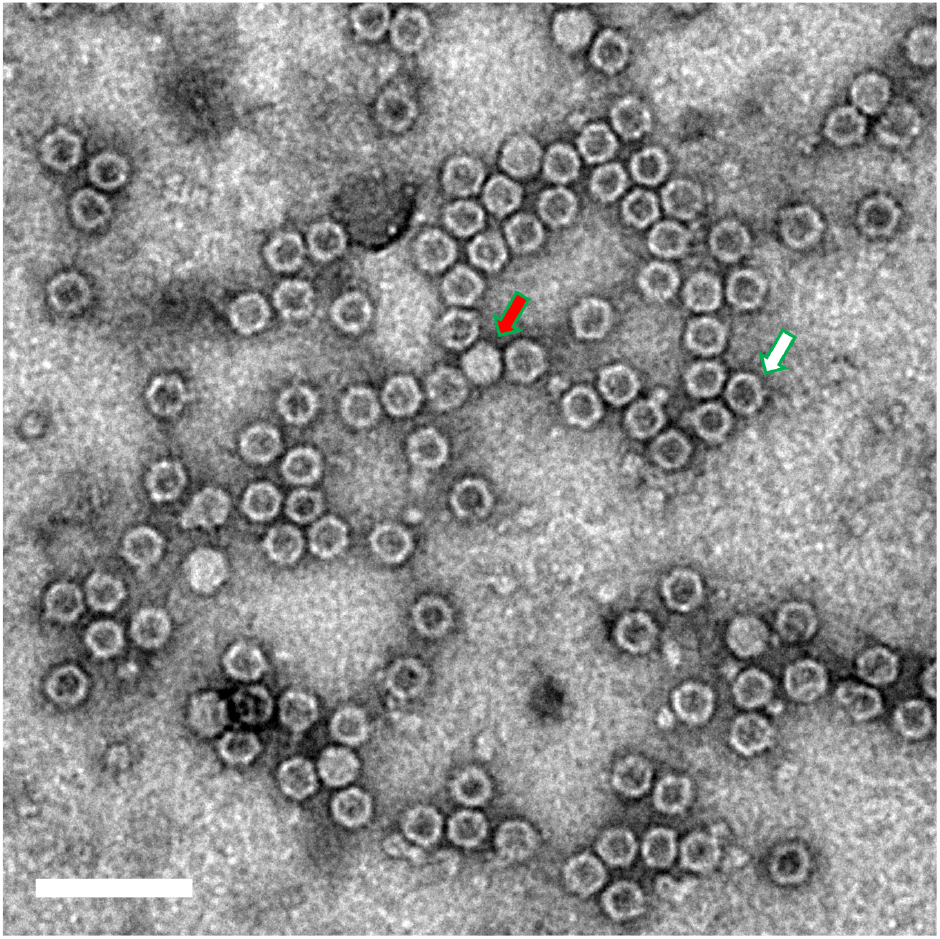
A negative-stain TEM image of the sample used in the MW-A UC analysis. Highlighted are a representative full capsid (red arrow) and a presumed empty capsid (white arrow). Scale bar: 100nm.

## Methods

### Design of MW-AUC experiments

We describe here how the features of each optical system are best exploited for multi-wavelength analytical ultracentrifugation experiments involving biopolymers, in particular with a focus on macromolecular interactions. We focus on the experimental design and describe how the spectral features of each analyte can be used to optimize the information obtained. We also discuss the algorithms used to analyze multi-wavelength data obtained from the Optima AUC since they differ from the earlier described procedure that is suitable only for the Cölfen optics [4].

Multi-wavelength AUC (MW-AUC) is a valuable method for investigating solution-based mixtures of interacting or non-interacting analytes, where each analyte contributes a unique chromophore. In addition to traditional single-wavelength methods, MW-AUC also characterizes the hydrodynamically separated molecules based on their spectral contributions, identifying free and complexed species they may form, as well as the stoichiometries of their complexes. This technique relies on the ability to spectrally separate the absorbing species present in a mixture. In order to successfully separate the spectral contributions from different analytes, several requirements need to be met. First, the mixing event should not induce a change of the analyte’s absorbance properties. For example, in the case of complex formation, the absorbance spectra of the interacting analytes should not red- or blue-shift, or change molar absorptivity. Second, the absorbance spectra of the pure analytes should be known, preferably in molar dimensions, such that molar stoichiometries can be derived from the analysis. Third, the absorbance spectra of the analytes need to be sufficiently orthogonal in order to be linearly separable. This requirement can be checked by calculating the angle *θ* between the molar extinction vectors *u* and *v* of two analytes to be spectrally separated:

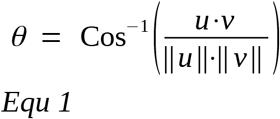

Theoretically, if the angle *θ* is larger than zero, the spectra can be separated. An angle of 90° indicates perfect orthogonality, but angles can be much smaller than 90° can be separated. The degree of success depends on the total signal available and the quality of the data. In general, the larger the angle *θ*, the better the chance the analytes can be spectrally separated. For analytes where the absorbance spectra show significant overlap (small *θ*), it is often helpful to expand the measured wavelength range. For example, when comparing the absorbance spectra of a typical protein and DNA, using just the typical 260 nm/280 nm absorbance pairs, *θ* is 27.8°, however, when considering the absorbance range between 230-300 nm, the angle increases to 42.7°, offering significantly improved resolution (also see Figure SI 1). The final requirement is that molar extinction profiles are within the same order of magnitude, ensuring that the observed signal is comparable between the different species. This can be a challenge when the molar extinction of a protein at 280 nm is much less than the molar extinction of an interacting nucleic acid at 260 nm. In such cases, mixtures quickly reach the dynamic range of the detector without providing sufficient signal from the protein. A solution is to shift or expand the measured wavelength range.

For example, nucleic acids have a particularly strong extinction in the 250-260 nm region, which partially overlaps with the 280 nm absorbance band of aromatic amino acids. Hence, measuring 240-300 nm works well for characterizing protein-nucleic acid interactions when the proteins contain a large mole fraction of tryptophan and tyrosine, and the nucleic acids are short [7]. Systems with longer fragments of nucleic acids in a mixture with proteins containing a small mole fraction of tryptophan and tyrosine will be challenging in multi-wavelength experiments conducted in this wavelength range, because the relatively small molar absorbance from aromatic amino acids is overwhelmed by the absorbance from the nucleic acid, and the protein will be difficult to detect. In such cases, sufficient signal from the protein can be achieved by including wavelengths in the region between 215-240 nm, were the peptide bond absorbance provides significantly higher absorbance (see Figure SI 1). This equalizes the absorptivity between protein and nucleic acid and at the same time increases the orthogonality between the absorption profiles of protein and nucleic acid.

In all cases it is important to use a buffer system that does not absorb significantly in the measured wavelength range. Suitable buffer systems include phosphate- or low concentration optically pure TRIS-based buffers, and do not contain absorbing additives such as nucleotides, chelators, or reductants in order to minimize the background absorbance.

For the Beckman optics, it is beneficial to minimize the number of wavelengths scanned because each wavelength has to be measured sequentially. That reduces the number of scans available for each individual wavelength compared to the Cölfen optics, which scans all wavelengths in parallel. One approach to maximizing the orthogonality of the measured spectra, while minimizing the number of measured wavelengths, is to interpolate spectral regions in the absorbance spectrum that exhibit linear change and to measure only wavelengths required for a faithful interpolation of the spectrum. For example, in regions where the change in the spectrum is linear over multiple wavelengths, only the endpoints of the linear region need to be measured. This will reduce the number of measured wavelengths and the time required to complete the scan cycle, thereby increasing the total number of scans collected for each wavelength. Another trick for the Beckman optics is to choose a rotor speed that is optimally synchronized with the flash lamp timing, which decreases the elapsed time between successive scans. The timing delays between scans, as a function of rotor speed, are calculated in the UltraScan data acquisition module for the Optima AUC (see Figure SI 3), optimal rotor speeds include 14,600-14,900, 31,500-32,900, 45,800-50,900, and 59,600-60,000 RPM. In these ranges, scan times are 8 seconds/channel or less. Unfortunately it is not possible in the Optima to scan only one channel of a cell. Therefore, for multi-wavelength AUC experiments acquired with the Optima AUC, it is advisable to run a single cell containing two samples, one in each channel sector, because a reference channel is not required when using UltraScan. Importantly, experiments should always be measured in intensity mode to reduce stochastic noise contributions to the data [26].

### Identification of basis spectra

For reversible hetero-associating systems, AUC can separate free and complexed species based on differences in their hydrodynamic properties. Once hydrodynamically separated, optical deconvolution can identify the molar contribution of each interacting partner in a complex, and provides the stoichiometry of binding [4, 7]. Reliable interpretation of the stoichiometry requires that reliable and pure molar extinction coefficients are known for each analyte in the mixture contributing to the absorbance of the sample over the entire spectral range examined in a MW-AUC experiment. To obtain these molar extinction coefficient profiles, high-quality absorbance scans of each analyte are required. Depending on the spectral properties of the analyte, the dynamic range of the detector (0.1 – 0.9 OD) can be readily exceeded at some of the selected wavelengths when only a single analyte concentration is measured. For example, the molar extinction coefficient for a protein at 215 nm can easily exceed the extinction coefficient at 280 nm by 1-2 orders of magnitude when aromatic sidechains are sparse or absent in the protein sequence (e.g., histones, collagen). To address this challenge, multiple dilutions need to be measured in the spectrophotometer. This approach ensures that overlapping wavelength ranges for one or more dilutions fall within the dynamic range of the detector, yielding a reliable intrinsic extinction profile over the entire wavelength range. To obtain the intrinsic extinction spectrum of an analyte over the entire wavelength range, the extinction profile fitter in UltraScan [3] is used to globally fit multiple dilution spectra from the analyte to sums of Gaussian terms using the Levenberg-Marquardt non-linear least squares fitting algorithm [27, 28] (see Figure SI 4). The fitted model is normalized with a known molar extinction coefficient (typically at 280 nm for proteins), which can be retrieved directly from the UltraScan LIMS database and derived from the associated protein sequence based on the molar absorptivity of the amino acid composition at 280 nm. The global molar extinction profile is used downstream to decompose experimental MW-AUC data into molar concentrations of spectral constituents (discussed below).

If the buffer used to dissolve the analytes absorbs in the measured wavelength range, then all absorbance measurements of the analytes of interest should be performed in a spectrophotometer blanked against the buffer. Also, since all MW-AUC experiments should be performed in intensity mode, the absorbing buffer must be considered as a separate spectral species in the downstream MW decomposition. In order to obtain its absorbance spectrum, the buffer’s absorbance profile must be measured by blanking the spectrophotometer with distilled water. We recommend to use spectrophotometers with a 1 cm pathlength, fitted with quartz cuvettes. For all studies reported here we used a benchtop GENESYS™ 10S UV-Vis spectrophotometer from ThermoFisher.

For reversibly interacting systems, the thermodynamic binding isotherms are most reliably determined by measuring MW-AUC experiments of multiple titration points with different ratios of the interacting partners mixed together [7]. The spectral decomposition module in UltraScan is used to obtain the mixing ratio from each titration point. The absorbance spectrum of the titration mixture and the intrinsic molar extinction spectra for each distinct analyte in the mixture (the *basis* spectra) are loaded into the program. The program will determine the overlapping wavelengths available from each spectrum, and use this range to calculate the molar composition, providing residuals to the fit. The program also reports on the angle *θ* (see Equ 1) between each pair of basis spectra (see Figure SI 5). By monitoring *θ*, the program can also be used to optimize the wavelength selection to aid in the experimental design. If a hypochromic or hyperchromic shift occurs in the absorbance profile due to mixing, the fitting residuals will appear to be non-random, and providing feedback on the suitability of including selected wavelength ranges in the decomposition.

### Analysis of MW-AUC experiments

Due to the design differences between the two multi-wavelength optical systems, experimental data differ in their structure and need to be analyzed with different strategies. The Cölfen optics collect all wavelengths simultaneously and provide a complete spectrum for the entire wavelength range, which is determined by the diffraction grating used in the optics [2]. As a result, each radial observation in the scan simultaneously produces a complete wavelength scan, where all observations are collected at the *same time* for each wavelength, producing a 3D scan image (absorbance as a function of wavelength and radius, see Figure 5, left panel). This image can immediately be decomposed to obtain isolated optical signals for each separated analyte in the mixture [4] for each radial point in each scan. In the Beckman optics, multiple wavelengths are collected sequentially, which causes each scan to be collected at a slightly *different time*. The time difference observed between the first and last wavelength collected for a multi-wavelength scan depends on the rotor speed and the total number of wavelengths collected, and is calculated by UltraScan. The difference in time between individual scans at different wavelengths is not obviously apparent from visual inspection of the 3D data (see Figure 5, right panel), but must be addressed before spectral decomposition can be performed.

**Figure 5:**
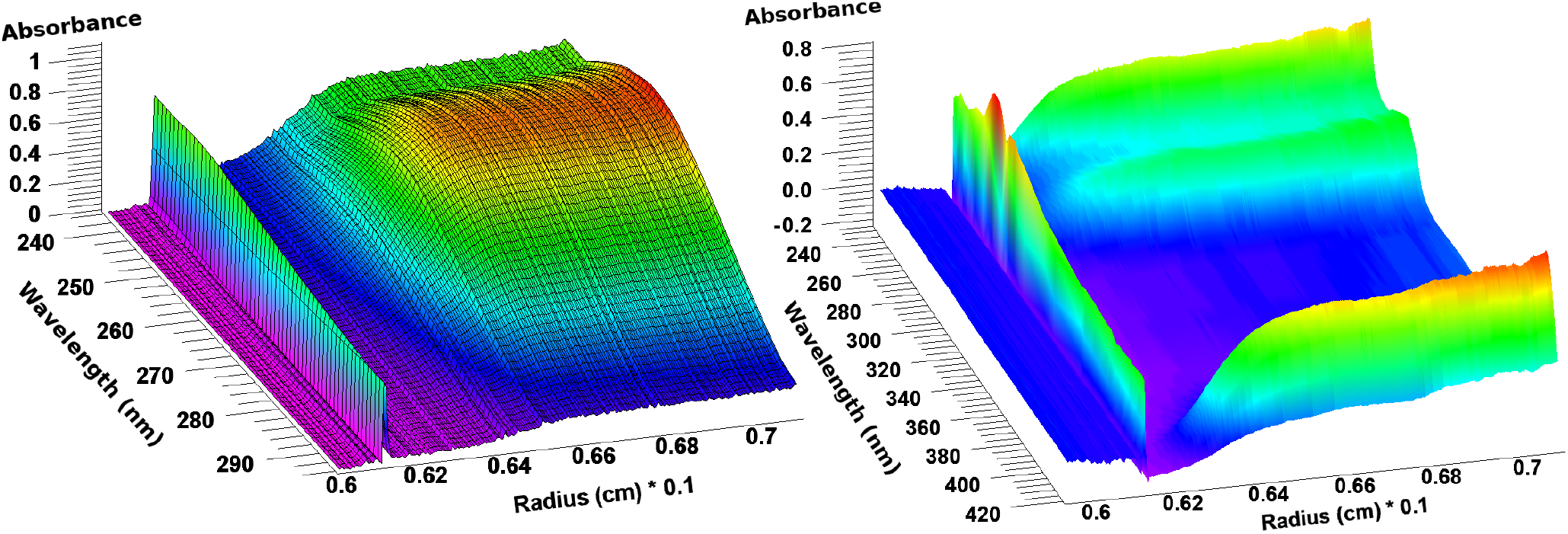
Multi-wavelength AUC data from a protein-RNA mixture acquired in the Cölfen optics (left) and a heme protein acquired from the Beckman optics (right). Only the Cölfen optics produce time-synchronous data, the displayed data from the Beckman optics contain wavelength data that are collected at different times, an issue that must be addressed before analysis. In both cases the meniscus is visible at the left, the sedimentation direction is to the right. The 410 nm heme peak is clearly visible in the right image.

For both optics, the analysis procedure before spectral decomposition is identical. The analysis starts by removing all systematic noise from each triple (a triple is a complete experimental dataset from a unique cell, channel, and wavelength) and fitting the boundary conditions (meniscus and bottom of cell). At this point, sedimentation velocity data from each triple are processed separately. The analysis proceeds through several refinement steps. In the first refinement step, a two-dimensional spectrum analysis (2DSA) [29] is performed with simultaneous time-invariant noise subtraction. In the Optima AUC, intensity data obtained from a photomultiplier tube contains significant time-invariant contributions, which must be removed first. This intensity variation is less of an issue with the linear CCD array used in the Cölfen optics, but the same step is still recommended to remove time invariant noise resulting from other sources, such as imperfections in the optical path or scratches in the cell windows. In the next step, a second refinement is performed with the 2DSA, adding time- and radially invariant noise correction, and fitting of the boundary conditions (meniscus and bottom of cell). On account of the mirror optics, both optical systems are essentially free of chromatic aberration [5]. However, in the Optima AUC, chromatic aberration in some instruments is large enough to require correction. This is handled in the UltraScan software by uploading a chromatic aberration profile into the LIMS database, which is applied to all data acquired from the Optima AUC during the data import stage. This process is further discussed by Stoutjesdyk et al. [30]. After chromatic aberration correction, the boundary condition fitting step only needs to be performed on a single wavelength from each channel, and the fitted positions is applied to the edit profiles of all other wavelengths in that dataset. For the boundary condition fit, it is recommended to select a wavelength which contains sufficient signal and low stochastic noise contributions. In the final refinement, an iterative 2DSA is performed with simultaneous time- and radially invariant noise correction for each triple [29]. The simultaneous processing of hundreds of triples for multiple channels and wavelengths is best performed on a supercomputer, where all triples can be analyzed in parallel and in batch mode [31]. Sedimentation and frictional ratio parameters for the 2DSA fits should be carefully adjusted to capture all hydrodynamic species in the sample. Fits for all triples should be inspected to ensure the fits result in random residuals for all scans and all wavelengths of a dataset using the Finite Element Model Viewer in UltraScan. At this point, all systematic noise contributions should be removed from the data, and the final 2DSA refinement can be expected to be an accurate representation of the underlying data. The analytes contained in the 2DSA models will faithfully reproduce the hydrodynamic profiles from the experimental data, and the random residuals in the fitted data should only represent the stochastic noise contributions and have Gaussian distributions.

### Generation of a synchronous time grid for Optima AUC data

Before multi-wavelength data are decomposed into spectral basis vectors, one additional step is required with data from the Beckman optics. All wavelength data from the same channel must be transposed onto a synchronous time grid to handle the time discrepancies incurred during sequential wavelength acquisition. This is accomplished by loading the iterative 2DSA models from each triple belonging to a single channel into the Optima multi-wavelength fit simulator (started from the “*Multiwavelength*” menu in UltraScan’s main menu). Using the 2DSA models, this module simulates the entire MW-AUC experiment, such that all sedimentation velocity experimental data from different wavelengths are now on a common and synchronous time grid. The synchronous time grid ensures that each scan from every wavelength has the same time stamp and can be used to obtain a reliable wavelength scan for each radial position. During the simulation of the synchronous time grid, the user can set all specifics of this simulation (rotor speed, meniscus position, run duration, number of scans) to match the settings of the original experiment (further described in SI-6). Partial concentrations of all analytes will be faithfully reproduced from the 2DSA models. Next, the simulations are uploaded to the LIMS database and edited to produce an equivalent MW-AUC experiment to the original experimental dataset. There is no requirement to add stochastic or other systematic noise to the data since all noise components have already been identified and subtracted from the data in earlier refinement steps. At this point, the data from the Cölfen optics and the Beckman optics are equivalent and can be further processed by the spectral decomposition module.

### Spectral decomposition of MW-AUC data

Spectral decomposition of MW-AUC data resolves species with unique chromophores in a mixture by their absorbance properties. Data processed as described above result in a de-facto wavelength absorbance scan for every time point (a scan) and radial position in the experiment. This wavelength scan can be decomposed into its basis absorbance spectra as described earlier and shown in SI 5. The decomposition is accomplished by using the non-negatively constrained least squares algorithm (NNLS) developed by Lawson and Hansen [32]. It assures that only positive contributions, or zero, are generated during the decomposition. For each basis vector, a two-dimensional (2D) space-time sedimentation velocity dataset will be generated during this process. Together, all basis vectors solve the linear equation subject to the constraint *x_i_* > 0 (see Equ 2):

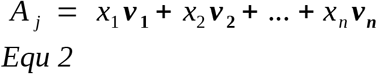

where *A_j_* is the absorbance wavelength scan at data point *j*, composed of spectral vectors ***V_i_*** with amplitudes *x_i_*. After processing all data in a MW-AUC dataset, the decomposition results in *n* traditional 2D sedimentation velocity experiments, each representing a separate, unique spectral species in the mixture. The decomposition is carried out by the UltraScan “*MWL Species Fit*” module from the “*Multiwavelength*” menu in the UltraScan main menu. This process is further detailed in SI-7. The resulting traditional 2D datasets (molar concentration as a function of radius and time) for each spectral component can be uploaded to the UltraScan LIMS system, edited, and analyzed by standard UltraScan procedures (2DSA [29], PCSA [33], GA [34, 35], van Holde – Weischet [36] or other methods available in UltraScan). There is no further need to fit the boundary conditions, remove systematic noise contributions, or perform a Monte Carlo noise analysis. Comparing spectrally separated hydrodynamic analyses will reveal both free and complexed species, where species with identical hydrodynamic parameters represent complexes. Importantly, integrating each spectral species found in a complex, the molar stoichiometry of the species in that complex is revealed, as long as the spectral basis vectors are expressed in terms of molar extinction coefficients[7].

### Preparations of fluorescent proteins

Hexahistidine-tagged fluorescent proteins (mPapaya1, Teal, and Ultramarine) were expressed in E. coli and purified by Ni-Sepharose chromatography according to methods described earlier [17, 18, 19].

### Preparations of AAV9 capsids

AAV9 capsids were produced in HEK293T/17 cells (ATCC, cat. CRL-11268) with the triple transient transfection method described before [37] and then purified with a commercial kit. Briefly, pUCmini-iCAP-AAV9 plasmid, pHelper plasmid, and a standard transgene cargo plasmid pAAV-CAG-GFP (Addgene #37825) were co-transfected to adherent HEK293T/17 cells at a mass ratio of 4:2:1. 3 days after transfection, the producer cells were lifted by adding 10mM EDTA to the media. After being spun down at 2000g for 10min, viral particles in the cell pellets were purified with AAVPro purification kit (Takara bio cat. 6675) following the manufacturer’s instructions. The concentration of genome-packaging capsids was quantified with real-time PCR (as described in Challis et al., 2019 [37]) using a pair of primers targeting the WPRE region. Particles with 5e12 packaged viral genomes were used for the AUC analysis.

## Discussion

The MW-AUC method extends the capabilities of an important biophysical characterization tool by adding a spectral characterization dimension to the hydrodynamic separation traditionally achieved by AUC. As is documented in three representative examples here, distinct advantages are realized in the resolution and information content for the study of heterogeneous systems when multiple analytes with unique chromophores are present in mixtures. This capability provides new avenues for the solution-based investigation of complex, interacting systems by providing higher resolution details about composition, binding strength, and stoichiometry of interaction than could be achieved with traditional AUC approaches. New instrumentation available in the form of the Cölfen and Beckman optical systems, as well as software advances in the UltraScan software, contribute to the advances reported here, and provide convenient access to this technology.

## Acknowledgments

This work was supported by the Canada 150 Research Chairs program (C150-2017-00015), the Canada Foundation for Innovation (CFI-37589), the National Institutes of Health (1R01GM120600) and the Canadian Natural Science and Engineering Research Council (DG-RGPIN-2019-05637). UltraScan supercomputer calculations were supported through NSF/XSEDE grant TG-MCB070039N, and University of Texas grant TG457201. Computational resources and support from the University of Montana’s Griz Shared Computing Cluster (GSCC) contributed to this research (all grants awareded to BD). The Canadian Natural Science and Engineering Research Council supports AH through a scholarship grant, and UK through RGPIN-2020-04965. The AAV work was supported by National Institutes of Health grant UF1MH128336 (to VG). Plasmids encoding fluorescent proteins were kindly provided by Dr. Robert Cambpell, University of Alberta.

## Author contributions

AH performed and analyzed all AUC experiments, and edited the manuscript. GEG, AS, and MK contributed to the multi-wavelength analysis modules software development in UltraScan. JH prepared and contributed the oil seed protein samples, SKS and UK prepared and contributed the fluorescent proteins, and VG and XD prepared and contributed the AAV samples.

## Supplemental Information

**SI 1:**
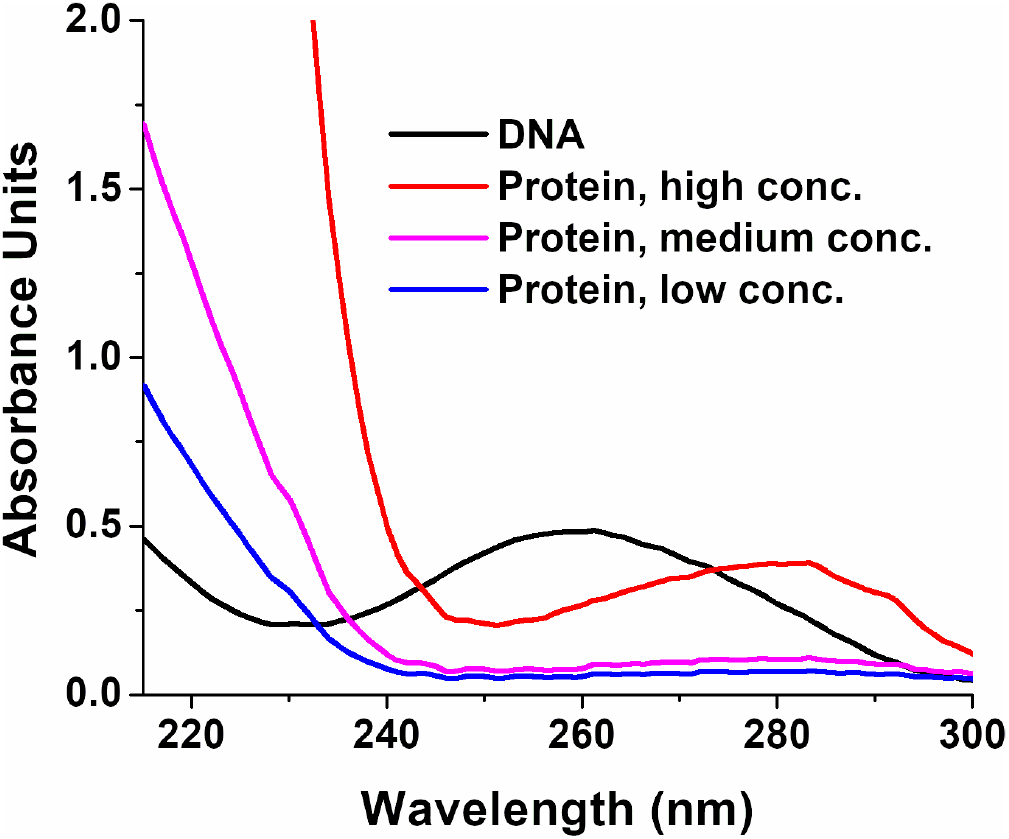
Absorption spectra for DNA and protein. Absorbance profiles for DNA (black) and protein (Bovine serum albumin, red: high concentration, magenta: medium concentration, blue: low concentration) between 210-300 nm. Even low concentration proteins, or proteins without aromatic side chains, provide sufficient signal and spectral orthogonality when wavelengths between 210-240 nm are included due to the absorbance from the peptide bond in the protein’s backbone.

**SI 2:**
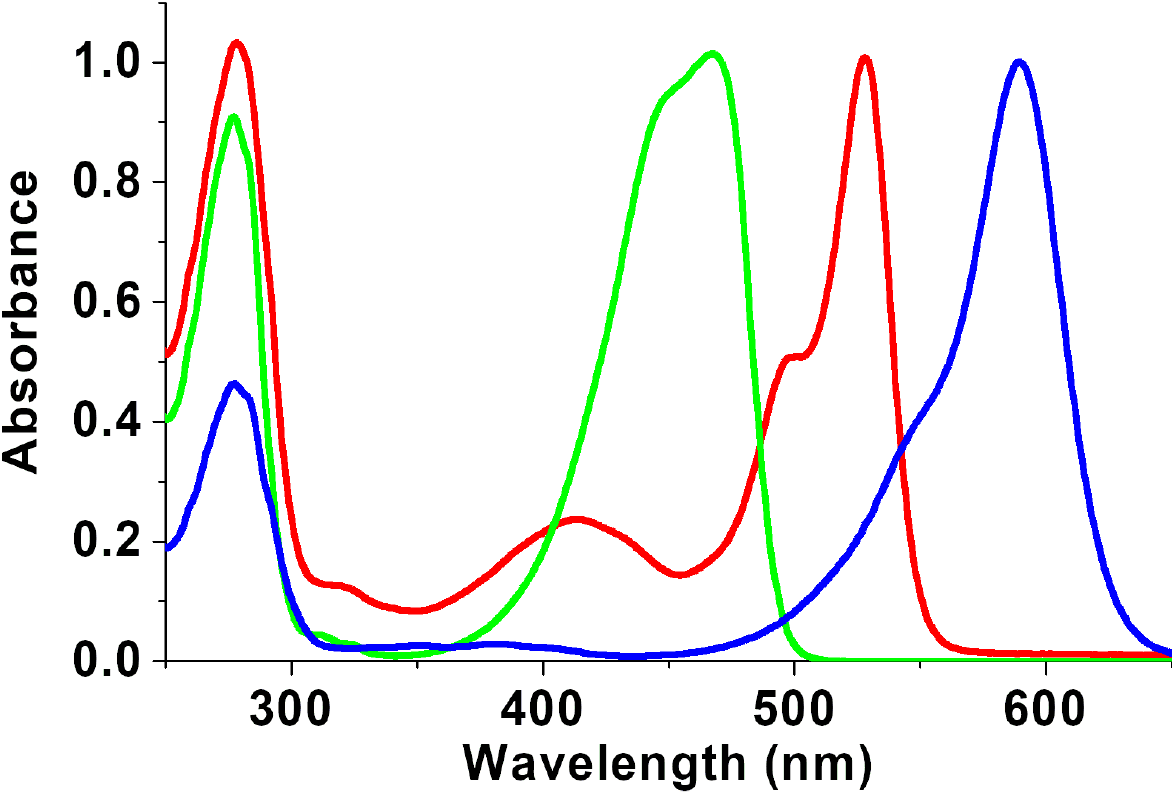
Normalized Absorbance Spectra for Fluorescent Proteins. Ultramarine (blue), mTFP1 (green), and mPapaya (red). While all fluorescent proteins share a peak at 280 nm due to tryptophan and tyrosine absorbance, the fluorescence excitation spectra in the visible region are markedly different and can be used to easily distinguish the spectra in a multi-wavelength AUC experiment. Proteins can be expressed as fusion proteins with fluorescent proteins to inherit unique excitation spectra from fluorescent proteins.

**SI 3:**
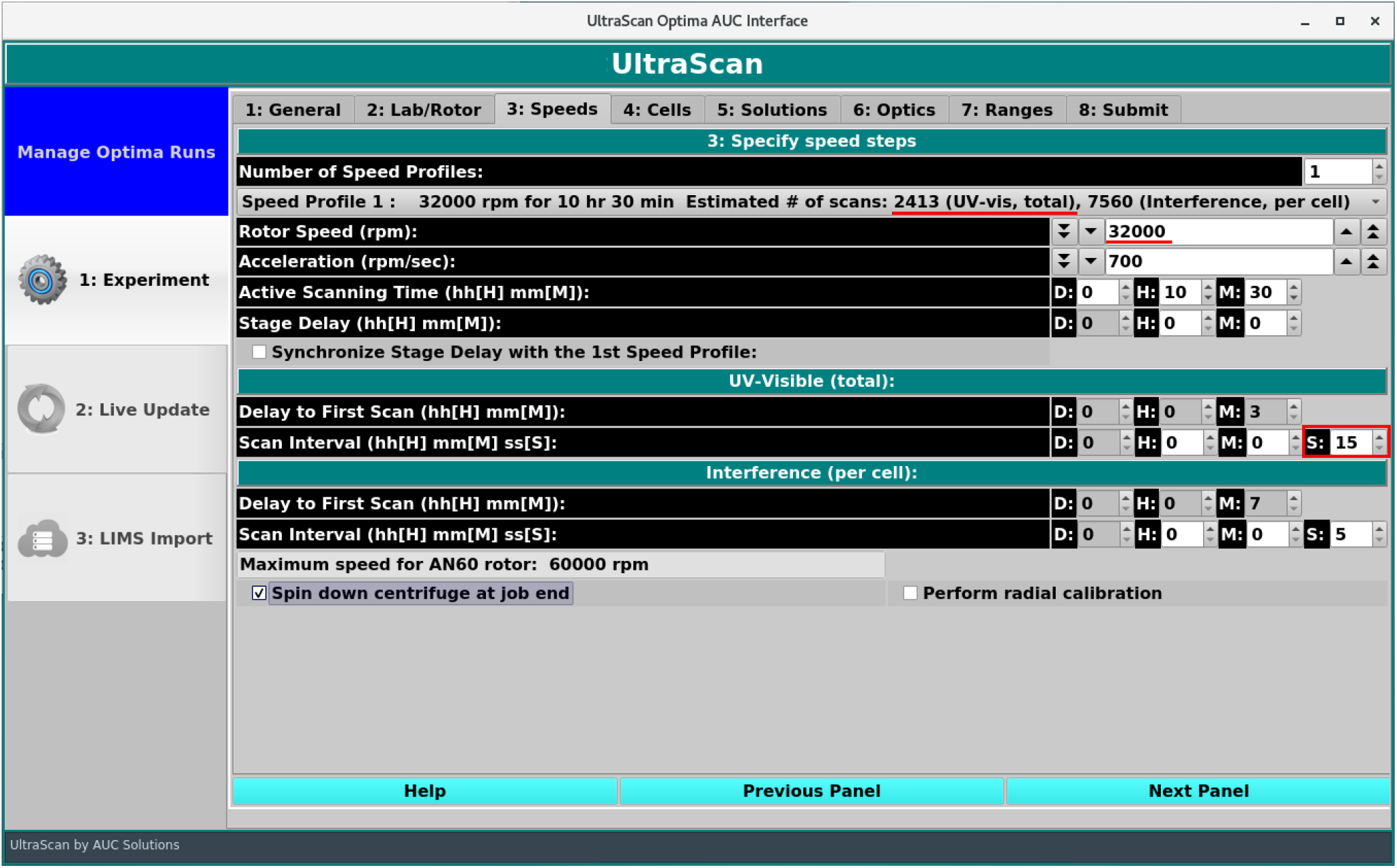
Speed selection control in the UltraScan data acquisition interface for the Optima AUC. Items marked in red relate the rotor speed selected for an experiment to the time interval in seconds between sequential scans, and the total number of scans that can be acquired over the user-selected data acquisition run time.

**SI 4:**
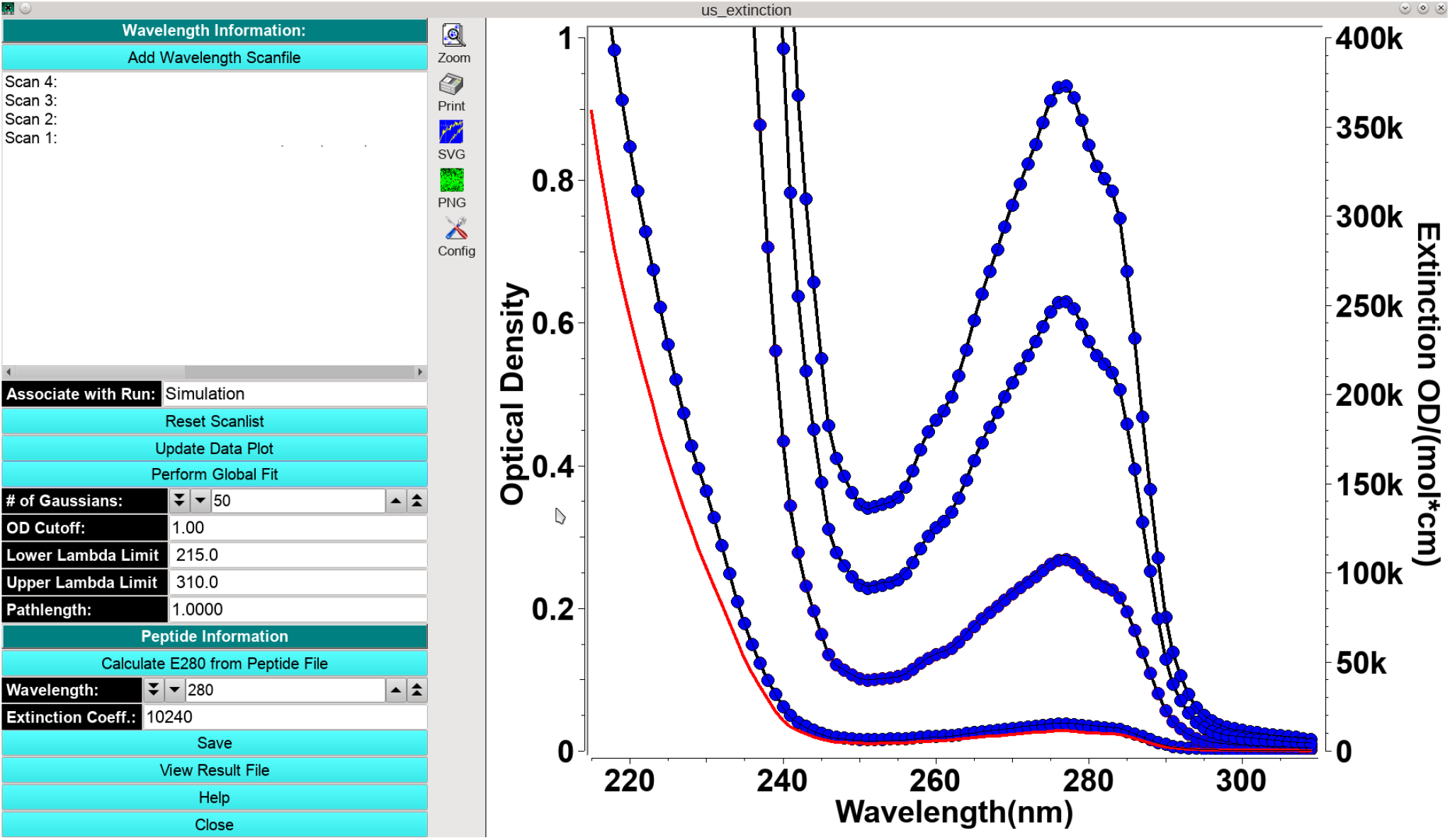
Global spectrum fitting interface. Experimental absorbance wavelength data from multiple analyte dilutions (blue dots, left y-axis) are fitted to a global molar extinction profile (red line, right y-axis). The global molar extinction profile model is precisely scaled to each concentration that was measured in a spectrophotometer (black lines).

**SI 5:**
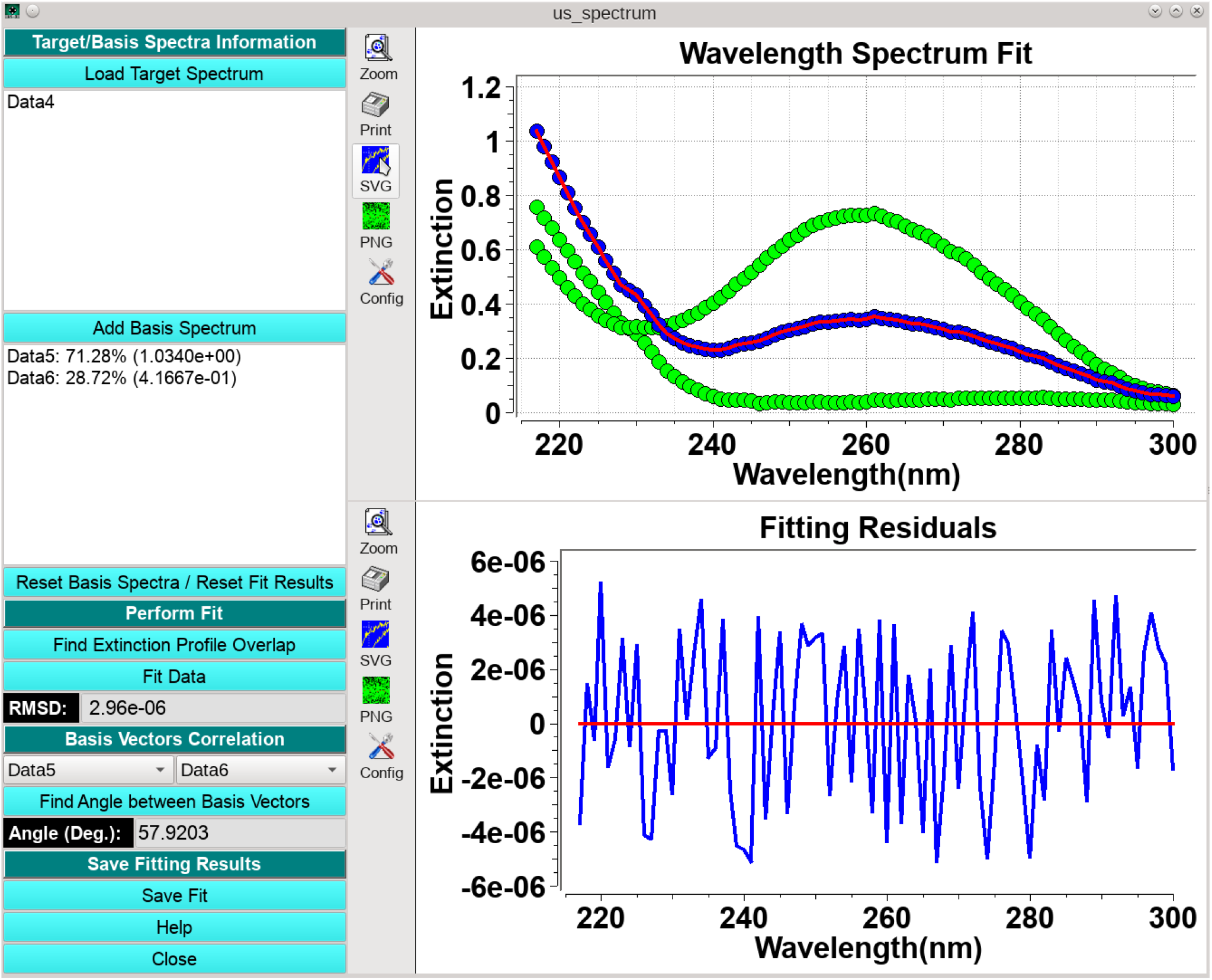
Spectrum decomposition utility in UltraScan. Absorbance scans of mixtures of spectrally diverse analytes (blue dots) are decomposed into their spectral basis vectors (green dots) and fitted to a linear combination (red line, above. Residuals of the fit are shown in the lower panel (blue line). Relative contribution of each basis is computed and displayed in the left panel. The angle between two basis vectors is displayed in the lower left.

**SI-6:**
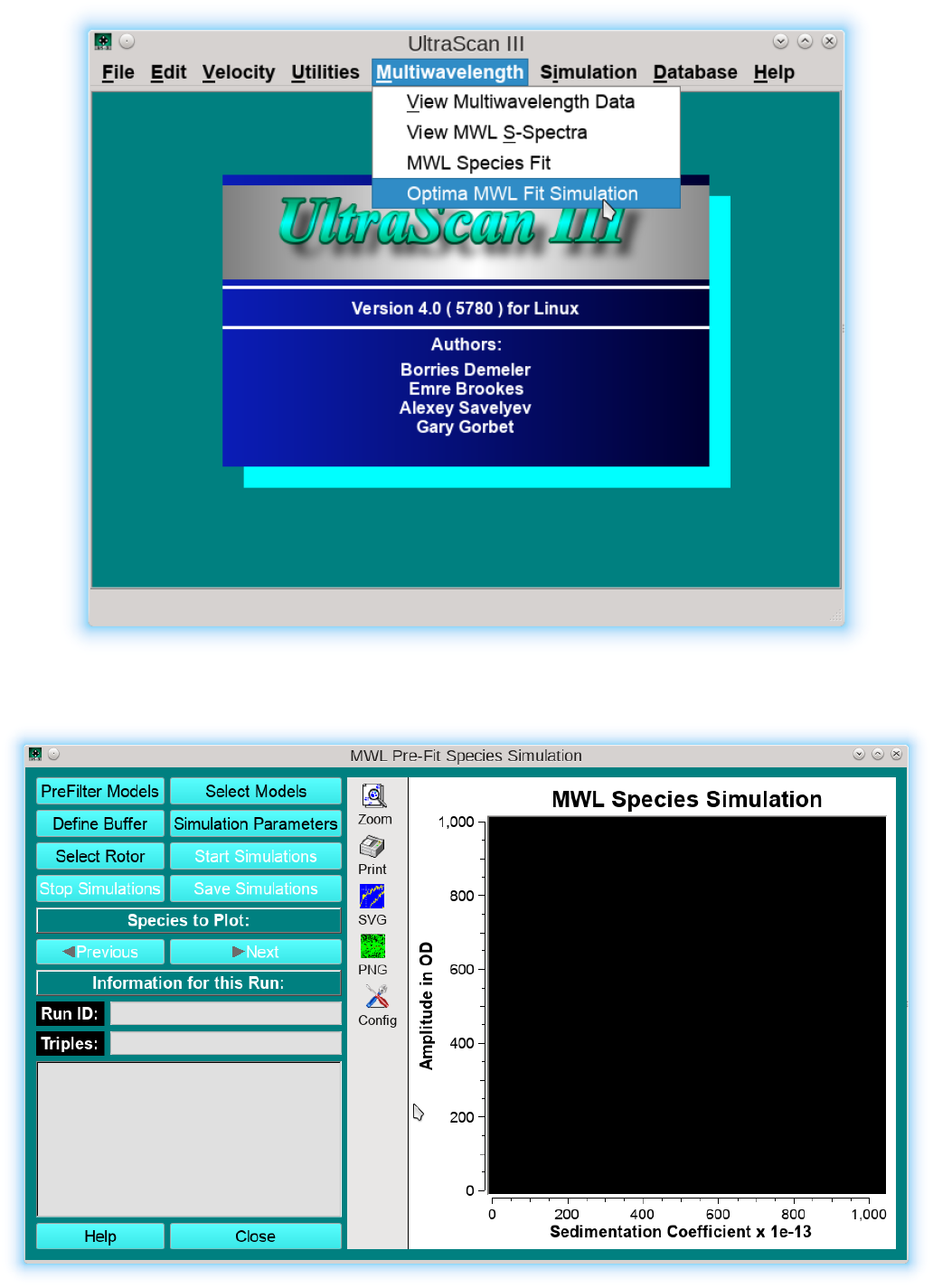
Step-by-Step instructions for the generation of time-synchronous multi-wavelength data from Optima AUC (Beckman Optics) intensity data. Step 1: Open the “Optima MWL Fit Simulation module from the main UltraScan menu:

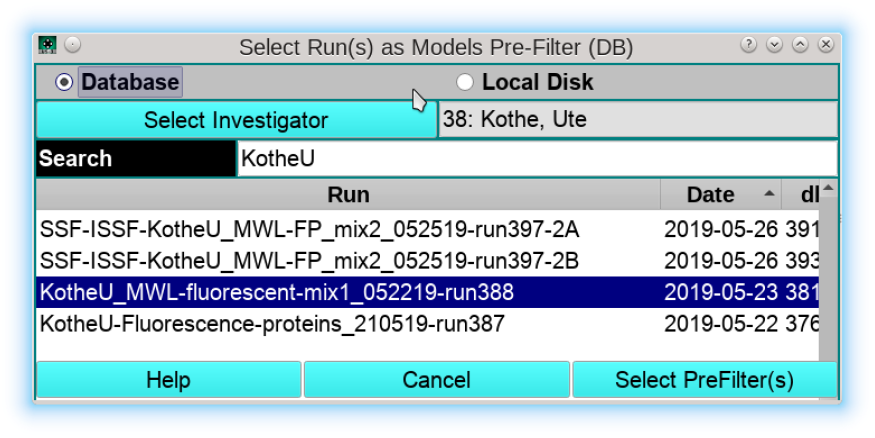
Step 2: Select a prefilter for the 2DSA-IT models to be simulated by clicking on “*PreFilter Models*”,and select the desired MW-AUC Optima AUC experiment:

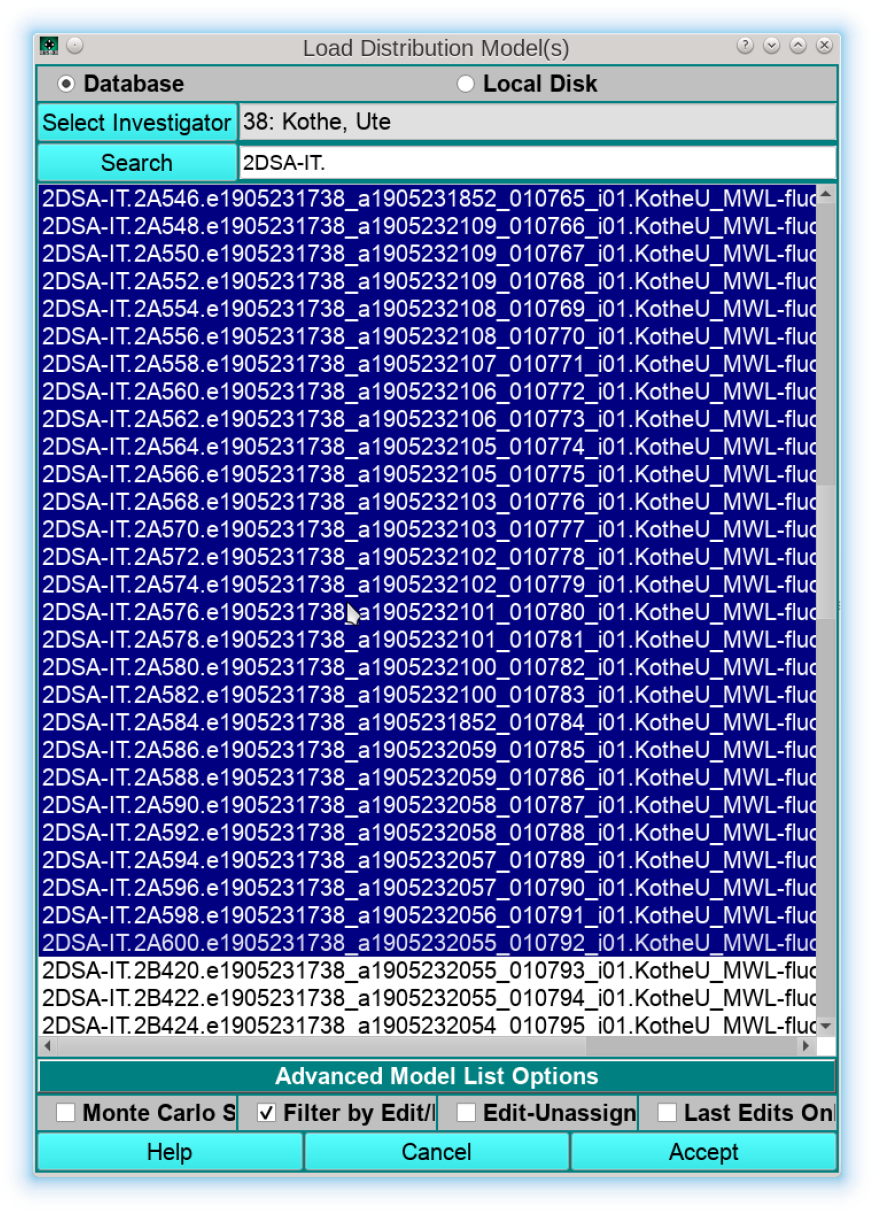
Step 3: Select all 2DSA-IT models for all wavelengths belonging to a single channel. By default, the program will display only 2DSA-IT models:

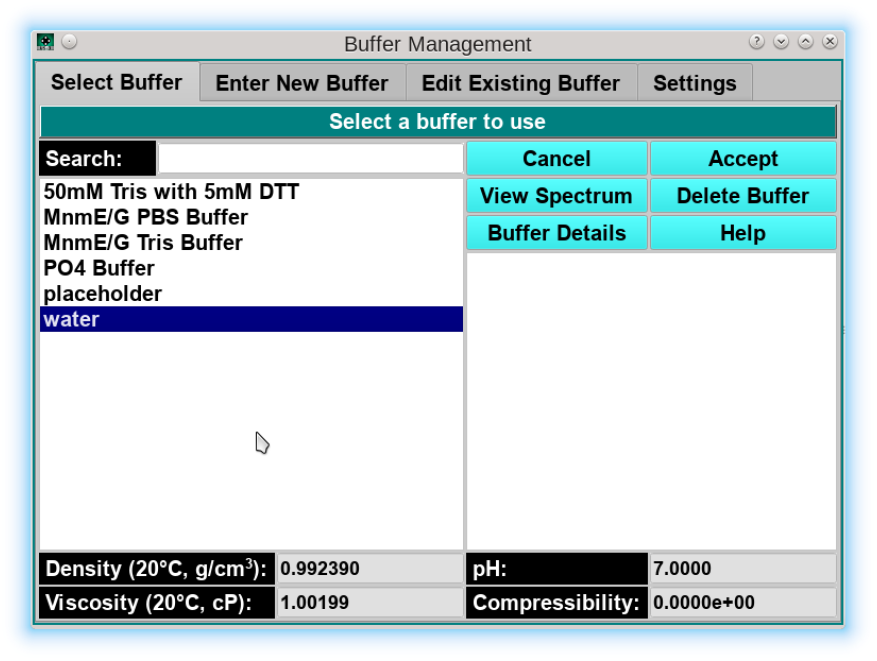
Step 4: Define a buffer by clicking on “*Define Buffer*”. Since buffer density and viscosity were already taken into account during the original 2DSA analysis, all 2DSA-IT models are already corrected for standard conditions, i.e., water at 20°C. Therefore, the user can pick water at 20°C for all subsequent analysis steps as a buffer:

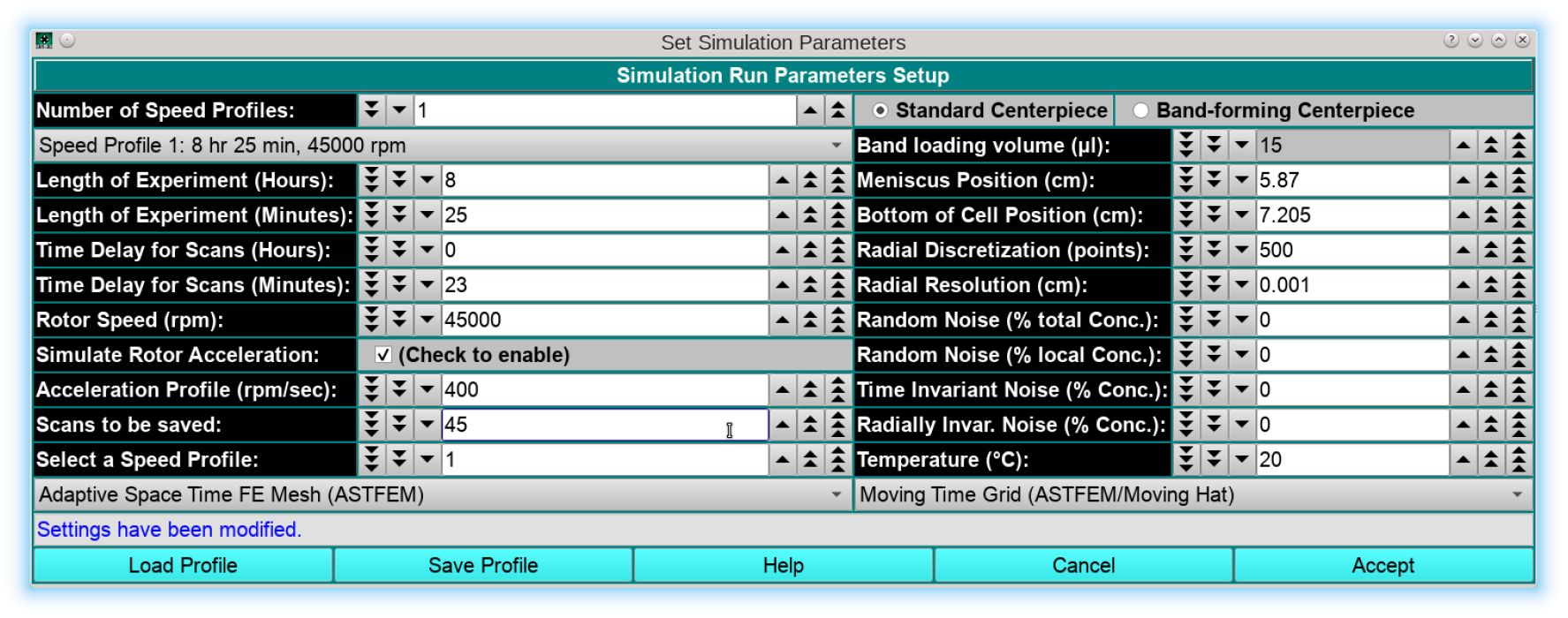
Step 5: In this step, the “*Simulation Parameters*” need to be defined. Ideally, these parameters should be identical to the experiment’s run parameters, including rotor speed, meniscus position (can be retrieved from the associated edit profile after the meniscus fit), rotor type and calibration profile, number of scans, run duration and scan delay should all be adjusted. It is important to note that UltraScan will report sedimentation and diffusion coefficients already corrected to standard conditions (20°C and water), so any simulations using the previously fitted 2DSA-IT models should use standard conditions:

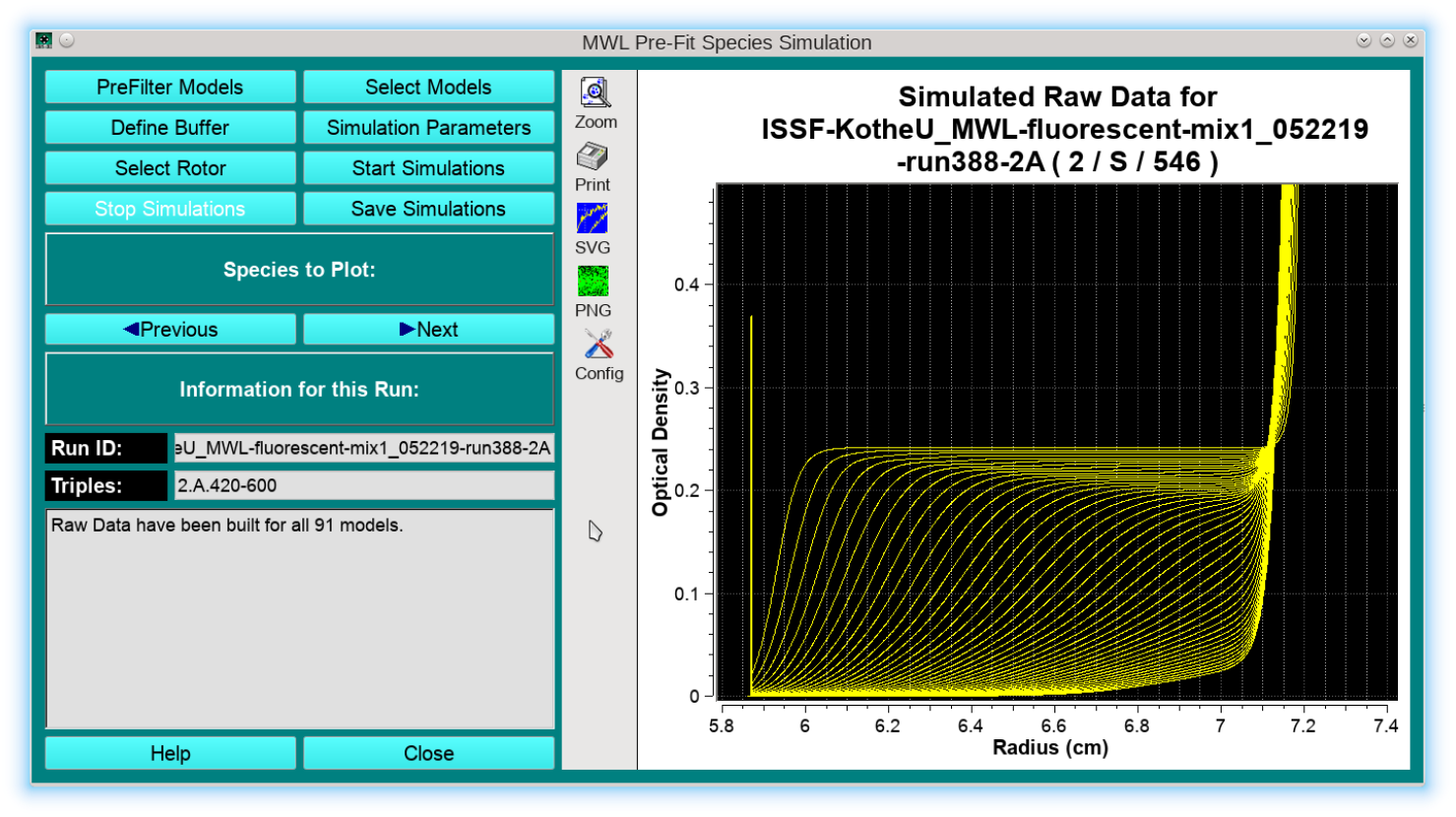
Step 6: Perform the simulation by clicking on “*Start Simulation*”. The program will re-simulate the fitted 2DSA-IT models with the experimental parameters defined in the simulation settings, generating a separate sedimentation velocity experiment for each wavelength. Now, each scan from each wavelength will be simulated for precisely the same time in all datasets. The simulated datasets will be shown in the graph windows, and a synthetic meniscus is generated as well to aid in downstream editing of these data. Clicking on “*Previous*” or “*Next*” allows the user to review each simulated dataset:

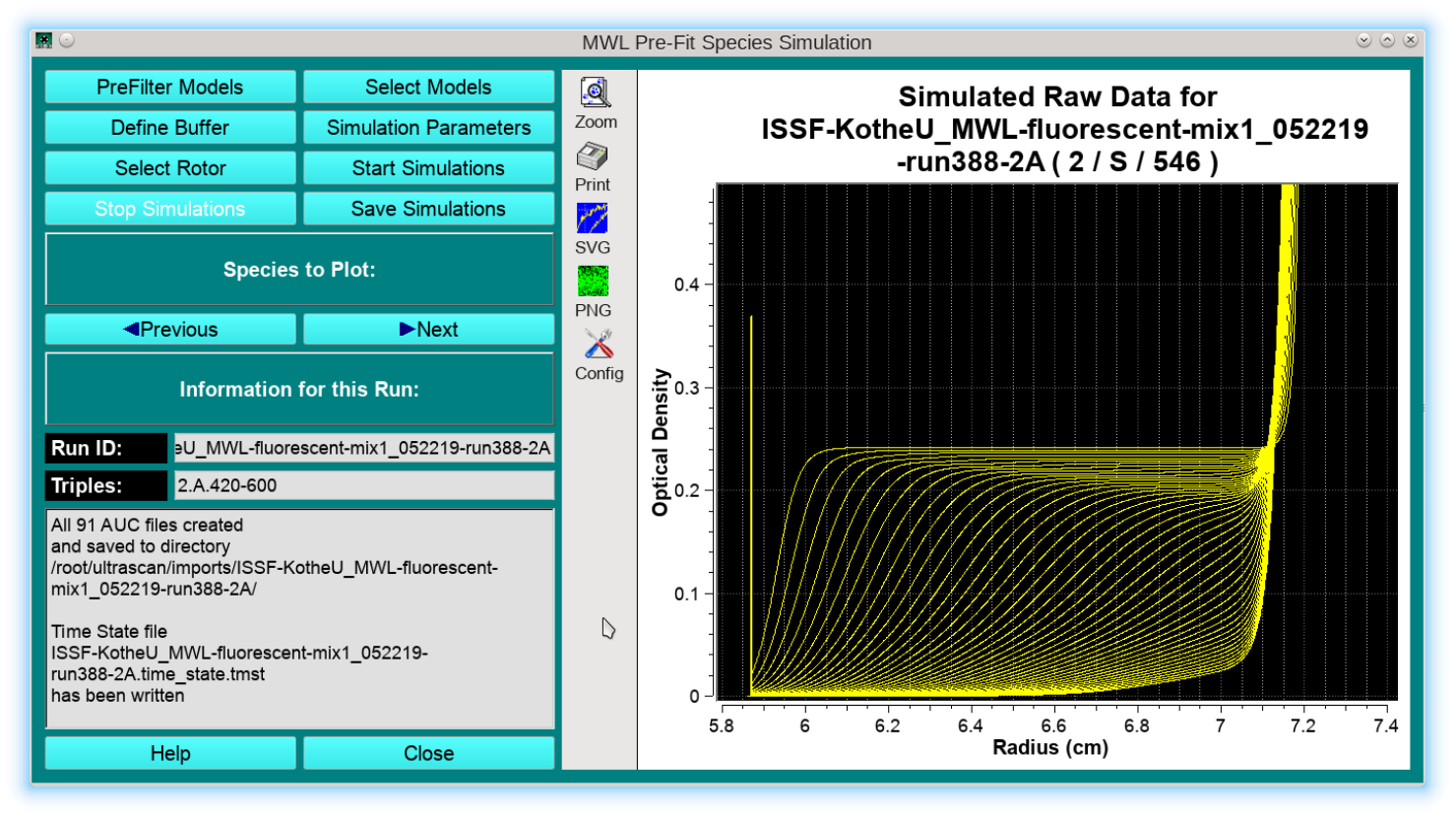
Step 7: Saving the data. Once the simulated data have been reviewed, the data can be saved by clicking on “*Save Simulations*”. Data will be written to the $HOME/ultrascan/imports directory into a subfolder that starts with prefix “*ISSF*-” (=initial simulated scan files): At this point, the ISSF data should be imported and edited like an ordinary MW-AUC velocity datasets.

**SI-7:**
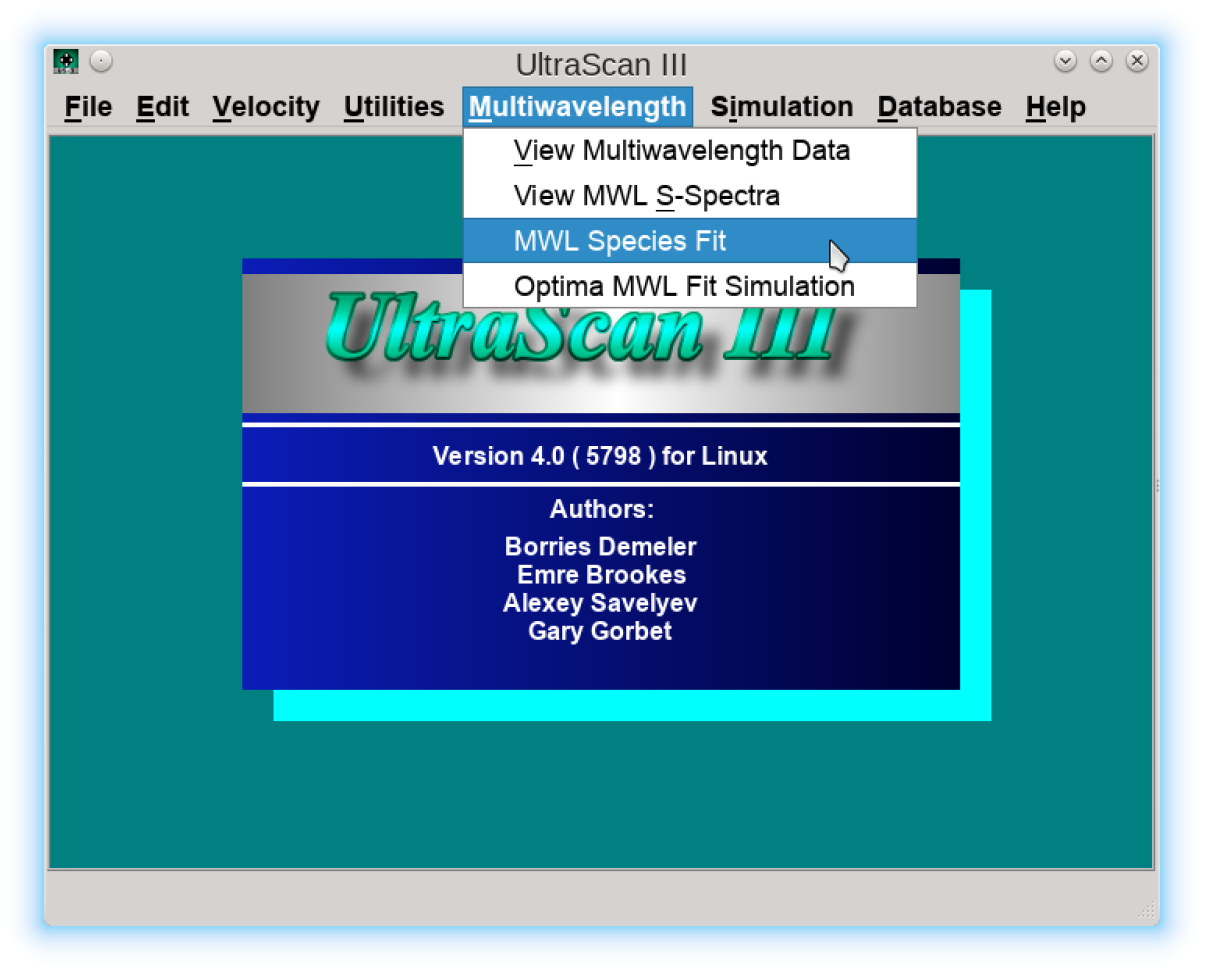
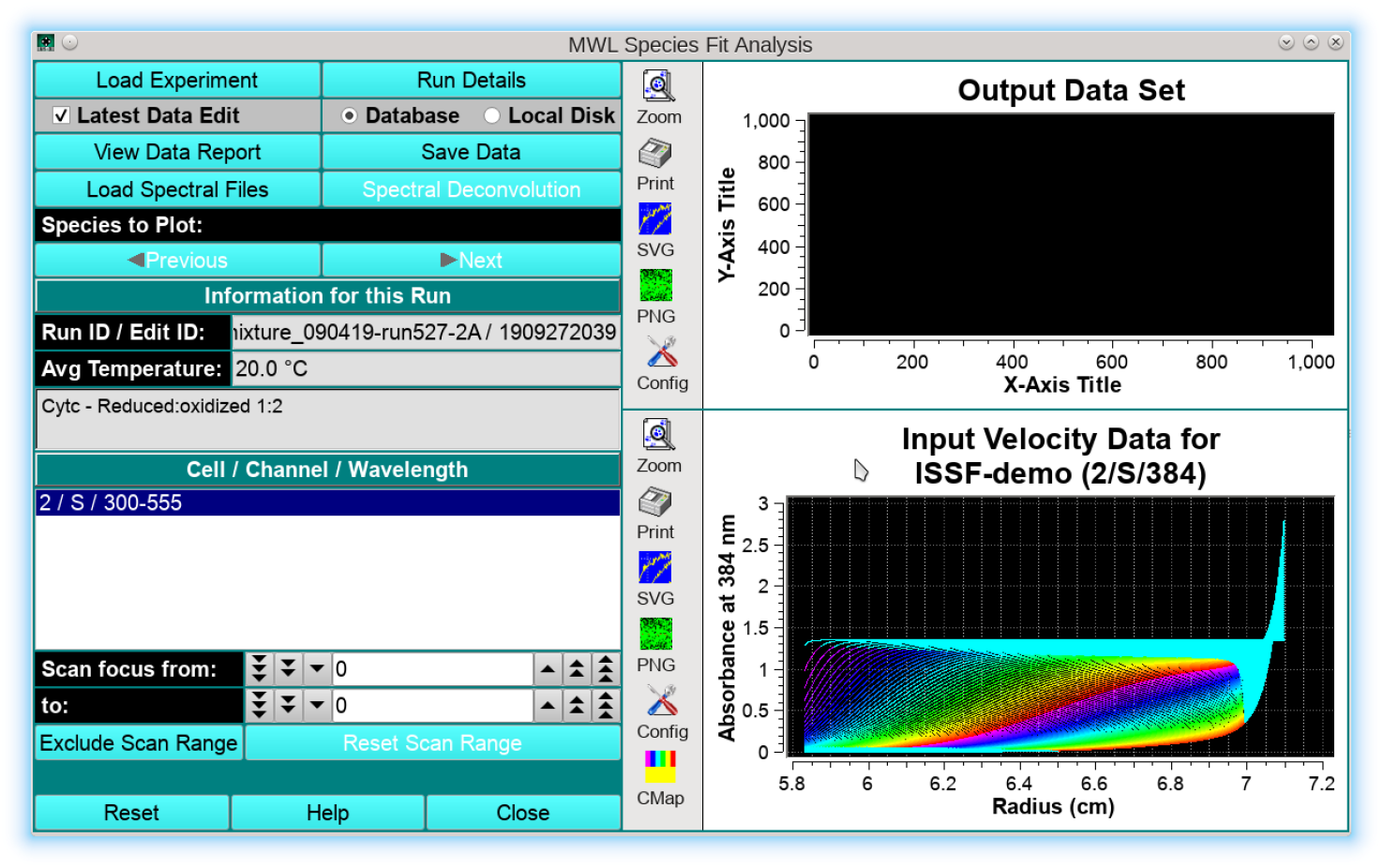
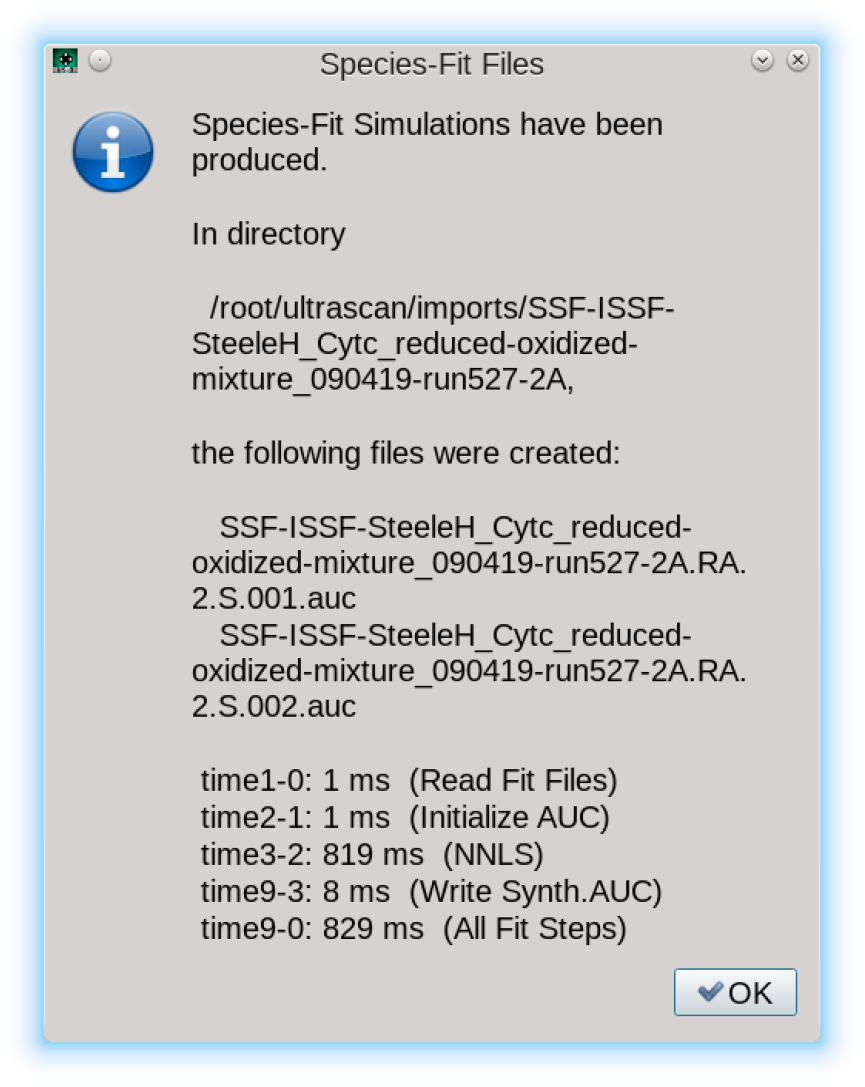
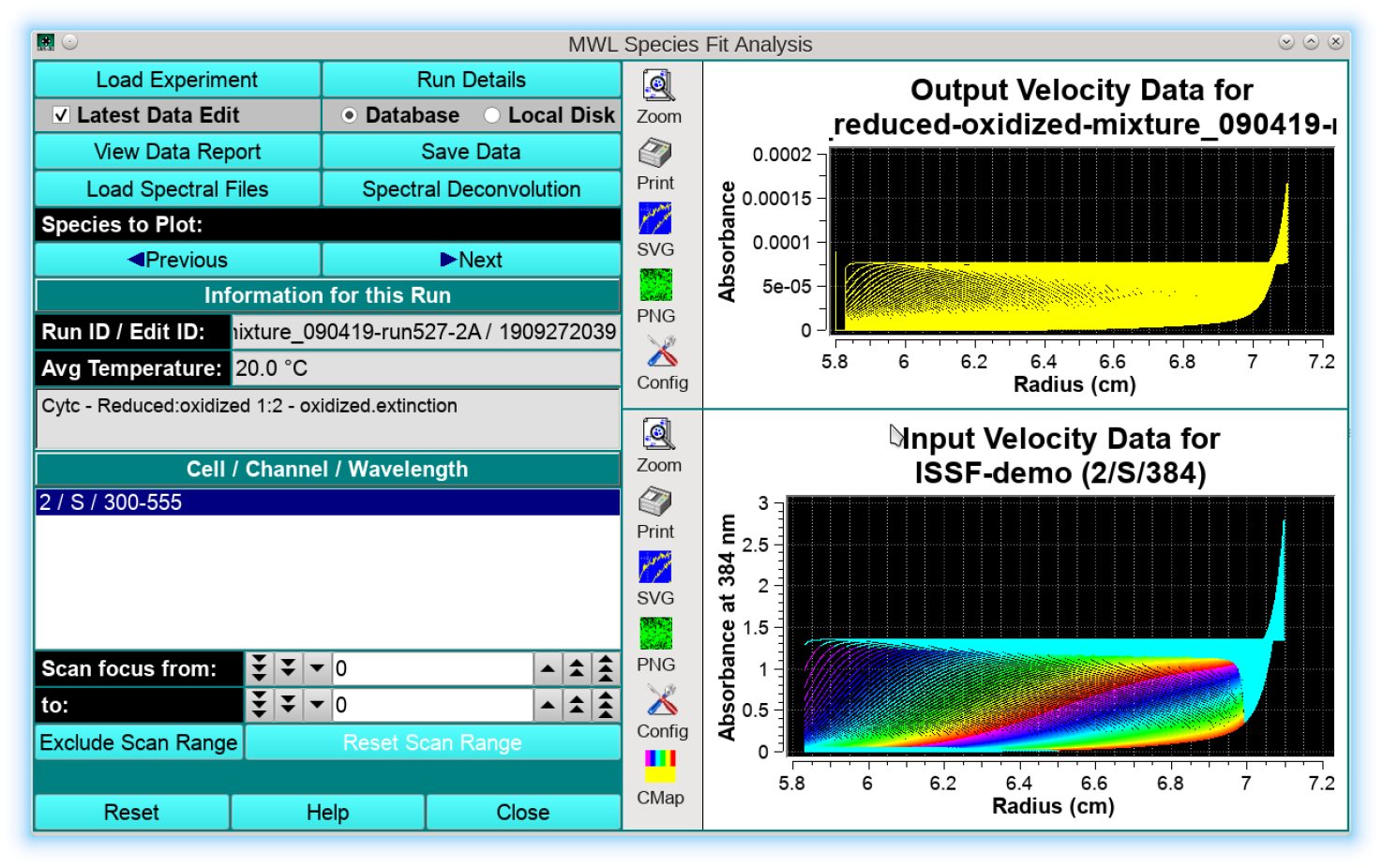
Step-by-Step instructions for the decomposition of time-synchronous multi-wavelength data. Decomposition of multi-wavelength sedimentation velocity experiments requires two or more extinction profiles for spectrally unique analytes present in a mixture, as well as a time-synchronous multi-wavelength dataset, either from the Cölfen optics or an ISSF dataset obtained after processing as described in SI-6, from the Beckman optics. The decomposition program is loaded from the main “*Multiwavelength*” menu entry by selecting “*MWL Species Fit*”: In the first step, the Cölfen optics or ISSF data are loaded into the program by clicking on “*Load Experiment*”: In the next step, the spectral bases vectors are loaded by clicking on “*Load Spectral Files*”. A minimum of two spectral bases need to be loaded. Once they are loaded, the “Spectral Deconvolution” button becomes active and should be clicked to start the deconvolution into separate datasets. The progress is reported in a dialog: Clicking on “*OK*” will reveal the deconvoluted datasets in the upper panel and activates the “*Previous*”and “Next” buttons to switch between datasets, and activates “*Save Data*” to save the results: The saved data need to be imported into the UltraScan LIMS server, edited and then they can be analyzed without any further meniscus or noise processing.

